# Glutathione Oxidation in Cerebrospinal Fluid as a Biomarker of Oxidative Stress in Amyotrophic Lateral Sclerosis

**DOI:** 10.1101/2024.07.01.601162

**Authors:** Trong Khoa Pham, Nick Verber, Martin R Turner, Andrea Malaspina, Mark O. Collins, Richard J. Mead, Pamela J. Shaw

## Abstract

**Background:** Oxidative stress is a key feature of several neurodegenerative diseases, including Amyotrophic Lateral Sclerosis (ALS). Identification of reliable biomarkers of oxidative stress would be beneficial for drug-target engagement studies.

**Methods:** We performed unbiased quantitative mass spectrometry (MS)-based analysis to measure changes in protein abundance and oxidation in cerebrospinal fluid (CSF) from a cohort of ALS patients and healthy controls at two time points (approximately four months apart) to capture disease progression. In addition, we developed a sensitive and targeted quantitative MS method to measure glutathione oxidation state in the same sets of CSF samples.

**Results:** Proteomic analysis of CSF revealed statistically significant changes in the abundance of several proteins, including CHIT1, CHI3L1, CHI3L2 and COL18A1 in ALS patients compared to healthy controls at both time points. Several sites of protein oxidation were significantly altered in ALS compared to healthy controls, and total levels of reversible protein oxidation were elevated in ALS patients. Given that glutathione oxidation could be a useful biomarker of oxidative stress, we also measured glutathione and its oxidation state in CSF in the same cohorts of samples. Total GSH (tGSH), GSSG levels and the GSSG/GSH ratio were significantly higher in the ALS than in the healthy control group for both time points. For the first visit, fold changes of tGSH, GSSG, and GSSG/GSH ratio in ALS compared to HC were 1.33 (p = 0.0215), 1.54 (p = 0.0041) and 1.80 (p = 0.0454), respectively. For the second visit, these values were 1.50 (p = 0.0143), 2.00 (p = 0.0018) and 2.14 (p = 0.0120), respectively. Furthermore, we found positive correlations between disease duration until the first visit and total glutathione (tGSH), GSSG and GSSG/GSH ratio. Finally, there was a strong positive correlation between the total intensity of reversibly oxidised proteins and the ratio of GSSG/GSH in ALS patients at both visits.

**Conclusion:** We propose that measuring levels of glutathione oxidation in CSF could act as a stratification biomarker to select ALS patients for antioxidant therapy and an approach to monitor the treatment response to therapeutic agents targeting oxidative stress.

## Introduction

Neurodegenerative diseases, such as ALS, are devastating conditions which present a distressing burden to those afflicted and society in general. Existing approaches have made few inroads towards the goal of slowing the progression of these diseases. We set out to develop biochemical biomarkers of target engagement (a direct measure of drug action) to measure the effects of therapies targeting oxidative stress, including edaravone and NRF2 (NF-E2-related factor 2) activators [1], which show great promise as disease-modifying agents for neurodegenerative diseases. Such ‘translational biomarkers’ provide a link between preclinical models and human clinical trials, facilitate dose selection, speed up clinical therapeutic development, provide early clinical proof-of-concept and are now seen as critical parameters for successful drug development [2]. In addition, molecular biomarkers of disease progression are needed to assess the efficacy of new treatments for patients with ALS. Since the brain is a difficult organ to access, changes in the abundance of proteins or peptides or their oxidation state can be measured in biofluids such as cerebrospinal fluid (CSF) [1].

Oxidative stress is considered a crucial factor directly or indirectly involved in several neurodegenerative diseases [3-5]. Many biomolecules are subject to oxidation, of which proteins and peptides are important candidates as biomarkers of oxidative stress in biofluids. Oxidative modifications can occur on amino acid residues such as methionine or cysteine (Cys), and depending on the level of oxidative stress, the oxidation of Cys can be either reversible or irreversible. Mass spectrometry (MS)-based approaches have been developed and applied for the large-scale detection and quantification of Cys oxidation in diverse biological systems [6]. However, the large-scale characterisation of Cys oxidation in neurodegenerative diseases is still limited, and the potential for Cys oxidation as a biomarker for disease progression and target engagement in ALS is unexplored.

Glutathione is a major antioxidant molecule which regulates the redox balance in the brain and is a promising therapeutic target for neurodegenerative disease treatment [7]. It has a tripeptide structure consisting of glutamate, cysteine and glycine, existing in neurons at concentrations ranging from 0.2 to 2 mM [8]. It regulates crucial cellular processes, including the metabolism of oestrogens, prostaglandins, leukotrienes, and xenobiotic drugs [8]. It also acts as an antioxidant agent, enzyme cofactor, the major redox buffer, a cysteine storage system, and a neuromodulator in the central nervous system (CNS) [9]. Reduced glutathione (GSH) is a major antioxidant, with its concentration in the brain parenchyma (2-3 mM) much higher than that in either blood or CSF [10. GSH acts as a crucial antioxidant agent in the brain, providing a defence against oxidative stress by directly reacting with reactive oxygen species (ROS) or participating in enzyme-catalysed redox reactions. The combination of low levels of defence mechanisms and a high level of ROS production in the CNS indicates that the CNS may be vulnerable to oxidative stress [10].

Glutathione exists in two forms: reduced glutathione (GSH) and oxidised glutathione (GSSG). Levels of GSH, GSSG and the GSSG/GSH ratio have been used to measure oxidative stress levels in many diseases, such as diabetes mellitus [11], cystic fibrosis [12], multiple sclerosis [13], mitochondrial disease [14] and Parkinson’s disease [15]. In the pathophysiology of ALS, oxidative stress is one of the key mechanisms which contribute to motor neuron injury and degeneration [16, 17]. Serum GSH is reported to be lower [18], and the GSSG/GSH ratio is higher [19] in patients with ALS compared to controls, suggesting an impaired capacity to buffer free radicals. Furthermore, analysis of plasma using Raman spectroscopy has implicated the metabolism of glutathione in disease progression [20]. A robust measure of the glutathione redox balance could be used to measure target engagement of antioxidant therapeutic approaches and define a patient sub-set that may benefit from novel disease-modifying treatments.

To identify potential biomarkers of disease progression and drug-target engagement, we developed mass spectrometry-based assays for the large-scale measurement of protein abundance, protein oxidation and glutathione oxidation in the same samples. We applied this workflow to analyse CSF samples from a cohort of ALS patients and healthy controls (HC). Proteomic studies have identified protein abundance changes associated with disease in cross-sectional studies, but few have assessed longitudinal changes. Therefore, we performed global and cysteine oxidative proteomics (oxi-proteomics) and targeted analysis of glutathione oxidation at two time points 4 months apart in ALS patients and HC. The proteomic analysis enabled the quantification of 699 proteins, and of these, 5 proteins showed significantly altered abundance in ALS CSF compared to HC for both consecutive visits. We quantified the oxidation of 1,219 cysteine sites on 287 proteins and found that the oxidation of several sites was significantly altered in ALS compared to healthy controls. Finally, we demonstrate that glutathione oxidation is significantly increased in ALS patients compared to controls. Notably, the oxidation level was positively correlated with disease duration. We propose that glutathione oxidation may be a useful biomarker of disease progression and has the potential to be used as a marker of drug-target engagement and therapeutic efficacy in clinical trials of drugs that modulate oxidative stress in the CNS.

## Results

### A workflow to measure protein abundance, oxidation and glutathione redox state from the same CSF samples

To maximise the use of valuable CSF samples, we created a workflow enabling the separation and analysis of protein and glutathione from the same samples (**Figure 1**). In a typical proteomic workflow to analyse CSF, immunodepletion is performed to remove abundant proteins that would otherwise preclude deep coverage of the proteome due to the high dynamic range of this biofluid. Given that the protein concentration of CSF is lower than plasma or serum, a concentration step using a centrifugation filter is often included prior to immunodepletion. This generates two fractions: a concentrated protein fraction for immunodepletion and a low molecular weight fraction that is typically discarded. We have developed a targeted mass spectrometry method that has the required sensitivity to detect glutathione in this lower molecular weight fraction, allowing measurement of protein abundance and oxidation as well as the redox state of glutathione, all from the same aliquot of CSF.

**Figure 1:**
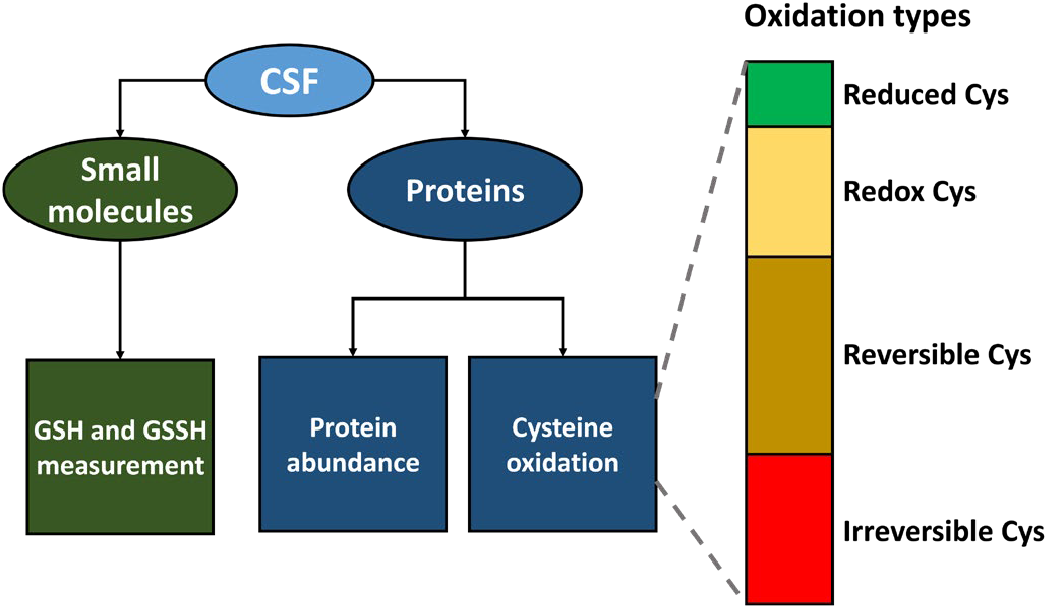
Oxidation types measured using our workflow. Reduced glutathione (GSH) and total reduced glutathione (tGSH) are measured using a targeted MS assay in a lower molecular fraction obtained from a protein concentration step. Oxidised glutathione (GSSG) levels are inferred from GSH and tGSH levels. Protein abundance and different types of protein oxidation on cysteine residues are measured by LC-MS/MS analysis. Reduced Cys: Cys residues identified in reduced form only; Redox Cys: Cys residues in both reduced and reversibly oxidised forms, the ratio of oxidised to reduced forms were used for comparisons between ALS patients and healthy controls groups; Reversible Cys: Cys residues identified in reversibly oxidised form only; Irreversible Cys: Cys residues identified in the irreversibly oxidised form only.

### Proteomic analysis of CSF from ALS patients and healthy controls

In this study, we employed a global redox proteomics method (Figure 1) to detect and measure different types of cysteine oxidation in CSF samples from ALS patients at two time points (visit 1, n = 24 and visit 2, n = 15) compared to HC (visit 1, n = 20 and visit 2, n = 11) (Table 1). This approach includes additional sample preparation steps and data analysis compared to a standard LC-MS/MS analysis to measure protein abundance. In this way, protein oxidation and protein abundance are measured simultaneously, with no additional analysis time. To maximise coverage of the CSF proteome, we used immunodepletion to capture the top 14 most abundant proteins and collected the flow-through for downstream analysis. To preserve the endogenous redox state of proteins in CSF, we employed differential alkylation of Cys residues before and after a reduction step. Proteins were then enzymatically digested, and the generated peptides were identified and quantified by tandem mass spectrometry. In total, 1,561 proteins were identified (ALS and HC groups) at a 1% FDR, and 699 proteins were quantified in 70% of samples (Supplementary Table S1). We used multiple ANOVA tests to identify differentially expressed proteins between the ALS and HC groups and between two consecutive clinic visits by ALS patients to determine their disease progression (Table 2). Five proteins were significantly increased in abundance in ALS compared to HC in both visits, including Chitotriosidase-1 (CHIT1), Chitinase-like proteins (CHI3L1 and CHI3L2), Collagen (COL18A1) and low-affinity immunoglobulin gamma Fc region receptor III-A (FCGR3A). No significant change in the abundance of these proteins in ALS CSF between two consecutive visits was observed. Furthermore, nine and fourteen proteins were significantly more abundant in ALS at the first and second visits, respectively (Table 2).

**Table 1.**
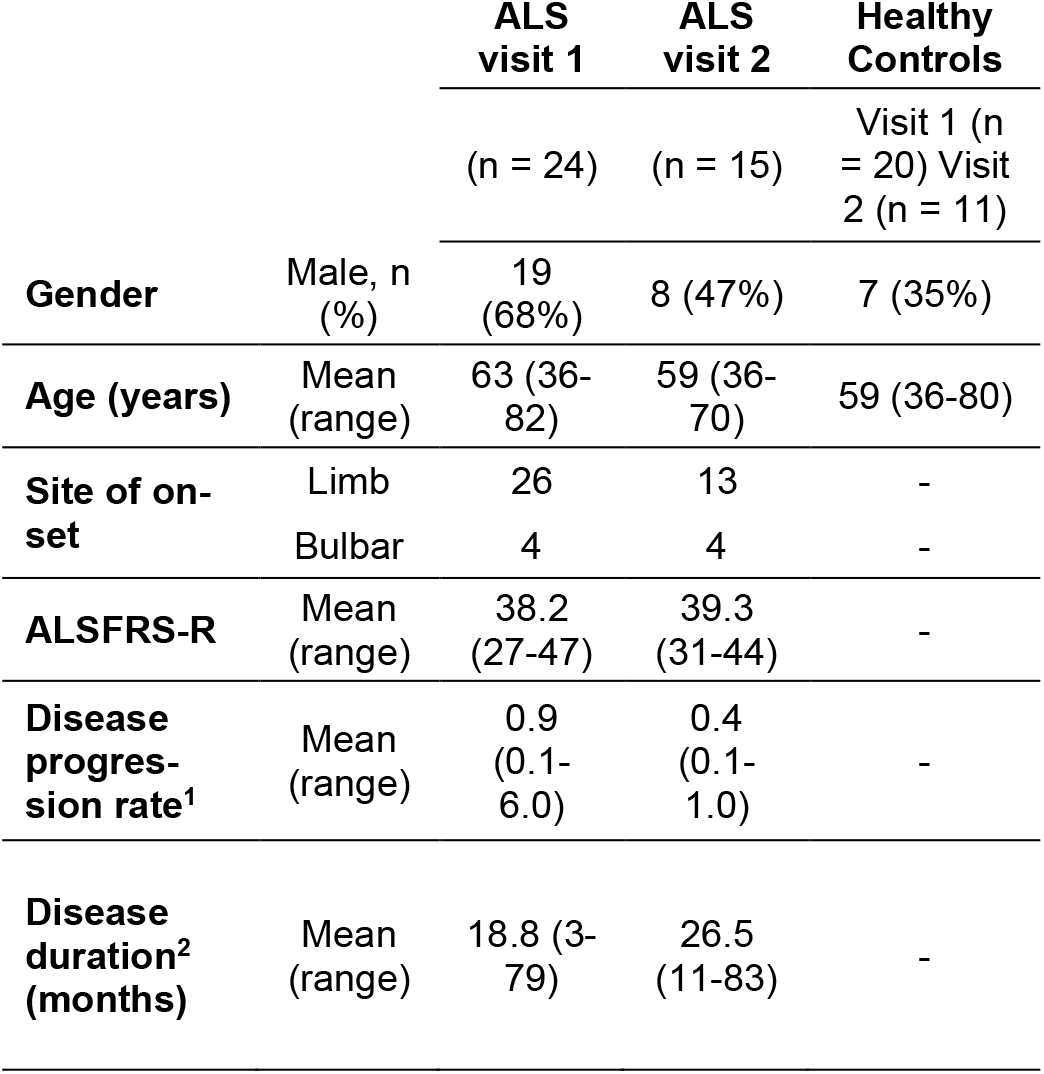
Subject demographics and clinical details. ^1^Disease progression rate = (48-ALSFRS-R)/disease duration (months). ^2^Disease duration = time from symptom onset until lumbar puncture

**Table 2.** Proteins with significantly altered abundance in ALS compared to HC groups in two consecutive visits. Bold values indicate statistically significant (Multiple sample ANOVA test) proteins in ALS vs HC (FDR 0.05). The full dataset is available in Supplementary Table S1.

| Gene | AC | Name | Log2 T-test difference 1st Visit | p Value 1st Visit | Log2 T-test difference 2nd Visit | p Value 2nd Visit |
| --- | --- | --- | --- | --- | --- | --- |
| CHI3L2 | Q15782 | Chitinase-3-like protein 2 | **1.49** | **1.20E-08** | **1.26** | **8.85E-06** |
| CHI3L1 | P36222 | Chitinase-3-like protein 1 | **0.70** | **1.80E-07** | **0.55** | **9.72E-04** |
| SEPINA3 | P01011 | Alpha-1-antichymotrypsin | **0.80** | **6.80E-05** | 0.31 | 5.95E-02 |
| CHIT1 | Q13231 | Chitotriosidase-1 | **2.88** | **1.30E-04** | **2.92** | **1.55E-03** |
| FCGR3A | P08637 | Low affinity immunoglobulin gamma Fc region receptor III-A | **0.52** | **1.70E-04** | **0.54** | **1.78E-02** |
| C7 | P10643 | Complement component C7 | **0.38** | **3.60E-04** | 0.14 | 2.51E-01 |
| MST1 | P26927 | Hepatocyte growth factor-like protein | **0.83** | **3.60E-04** | 0.26 | 2.88E-01 |
| CFD | P00746 | Complement factor D | **0.54** | **8.30E-04** | 0.08 | 5.54E-01 |
| CFH | P08603 | Complement factor H | **0.38** | **1.50E-03** | 0.06 | 6.90E-01 |
| C3 | P01024 | Complement C3 | **0.54** | **1.80E-03** | -0.01 | 9.53E-01 |
| MASP1 | P48740 | Mannan-binding lectin serine protease 1 | **0.52** | **2.60E-03** | 0.20 | 2.30E-01 |
| SERPING1 | P05155 | Plasma protease C1 inhibitor | **0.35** | **2.80E-03** | -0.01 | 9.46E-01 |
| MAN1B1 | Q9UKM7 | Endoplasmic reticulum mannosyl-oligosaccharide 1,2-alpha-mannosidase | -0.38 | 2.90E-03 | **0.18** | **5.85E-01** |
| COL18A1 | P39060 | Collagen alpha-1(XVIII) chain | **0.36** | **4.80E-03** | **0.29** | **5.22E-02** |
| NPC2 | P61916 | NPC intracellular cholesterol transporter 2 | 0.62 | 7.10E-03 | **0.69** | **1.73E-01** |
| EFEMP1 | Q12805 | EGF-containing fibulin-like extracellular matrix protein 1 | **0.52** | **1.60E-02** | 0.45 | 1.23E-02 |
| DMD | A0A087WV90 | Dystrophin | 1.85 | 1.10E-01 | **4.79** | **4.45E-04** |
| VASN | Q6EMK4 | Vasorin | 0.17 | 1.30E-01 | **0.37** | **8.41E-04** |
| SNED1 | Q8TER0 | Sushi, nidogen and EGF-like domain-containing protein 1 | 0.19 | 1.40E-01 | **0.41** | **1.01E-02** |
| CBR1 | P16152 | Carbonyl reductase [NADPH] 1 | -0.29 | 2.70E-01 | **0.61** | **7.62E-02** |
| CA11 | O75493 | Carbonic anhydrase-related protein 11 | 0.38 | 3.00E-01 | **0.91** | **2.49E-02** |
| VGF | O15240 | Neurosecretory protein VGF | -0.42 | 3.20E-01 | **1.36** | **2.47E-03** |
| PKM | P14618 | Pyruvate kinase PKM | -0.09 | 4.60E-01 | **0.78** | **2.69E-02** |
| GALNT10 | Q86SR1 | Polypeptide N-acetylgalactosaminyl-transferase 10 | 0.16 | 5.50E-01 | **1.19** | **3.63E-04** |
| PGK1 | P00558 | Phosphoglycerate kinase 1 | 0.06 | 6.30E-01 | **0.63** | **3.04E-02** |
| ECM2 | O94769 | Extracellular matrix protein 2 | 0.03 | 8.40E-01 | **0.28** | **2.27E-01** |
| PRG4 | Q92954 | Proteoglycan 4 | -0.01 | 9.80E-01 | **1.33** | **2.73E-03** |
| MAN2A1 | Q16706 | Alpha-mannosidase 2 | 0.00 | 9.90E-01 | **0.55** | **7.50E-03** |

We next investigated whether any protein oxidation biomarkers could be identified in the CSF of ALS patients. During the sample preparation before proteomic analysis, we performed differential alkylation of Cys residues to differentiate reduced and reversibly oxidised sites. Reduced Cys residues were alkylated using IAM, and reversibly oxidised Cys residues were reduced using DTT to generate newly formed sulfhydryl groups that were then alkylated using NEM. If a peptide containing Cys residues was labelled by both IAM and NEM, the intensities of reduced Cys (determined by IAM labelling) and oxidised Cys (by NEM) were compared to estimate the ratio of oxidised and reduced Cys. On the other hand, if a peptide containing Cys was only labelled by either IAM or NEM, it was considered to exist in only the reduced (IAM) or reversibly oxidised form (NEM).

The abundance of reduced and reversibly oxidised cysteine-containing peptides from different samples (ALS vs HC groups) was compared to estimate the ratio of cysteine oxidation. In addition, irreversibly oxidised cysteine forms such as dioxidation (8 sites) and tri-oxidation (55 sites) were identified. There were 47, 505, 531 and 63 cysteine sites detected as reduced only, redox (both reduced and oxidised), reversible (oxidised only) and irreversible, corresponding to 38, 143, 204 and 49 different proteins, respectively. Two proteins, dipeptidyl peptidase 2 (DPP7, Cys332 and Cys338) and complement factor B (B4E1Z4, Cys564), exhibited increased oxidation in ALS compared to HC in both visits. Brevican core protein (BCAN) containing two reversibly oxidised cysteines at positions Cys688 and Cys699 were significantly different in ALS compared to HC for both consecutive visits (Supplementary Table S2). In the ALS group, the cysteine oxidation status of some proteins was increased between two visits, but their oxidation abundances were not significant compared to the HC group (either the first or second visit). There were no differences in irreversibly oxidised cysteine status in ALS vs HC groups for both visits, and only protein Alpha-1-antichymotrypsin (SERPINA3C) had an increase in its tri-oxidised form in ALS vs HC for the second visit (Supplementary Table S2). However, there was a significant increase in the total intensity of reversibly oxidised Cys (the sum of all oxidised peptide intensities)) between the ALS patient group and HC group at the first visit, as shown in Figure 2 (1.23 fold, p = 0.0405), but this increase was not statistically significant at the second visit (1.19 fold, p = 0.3039).

**Figure 2:**
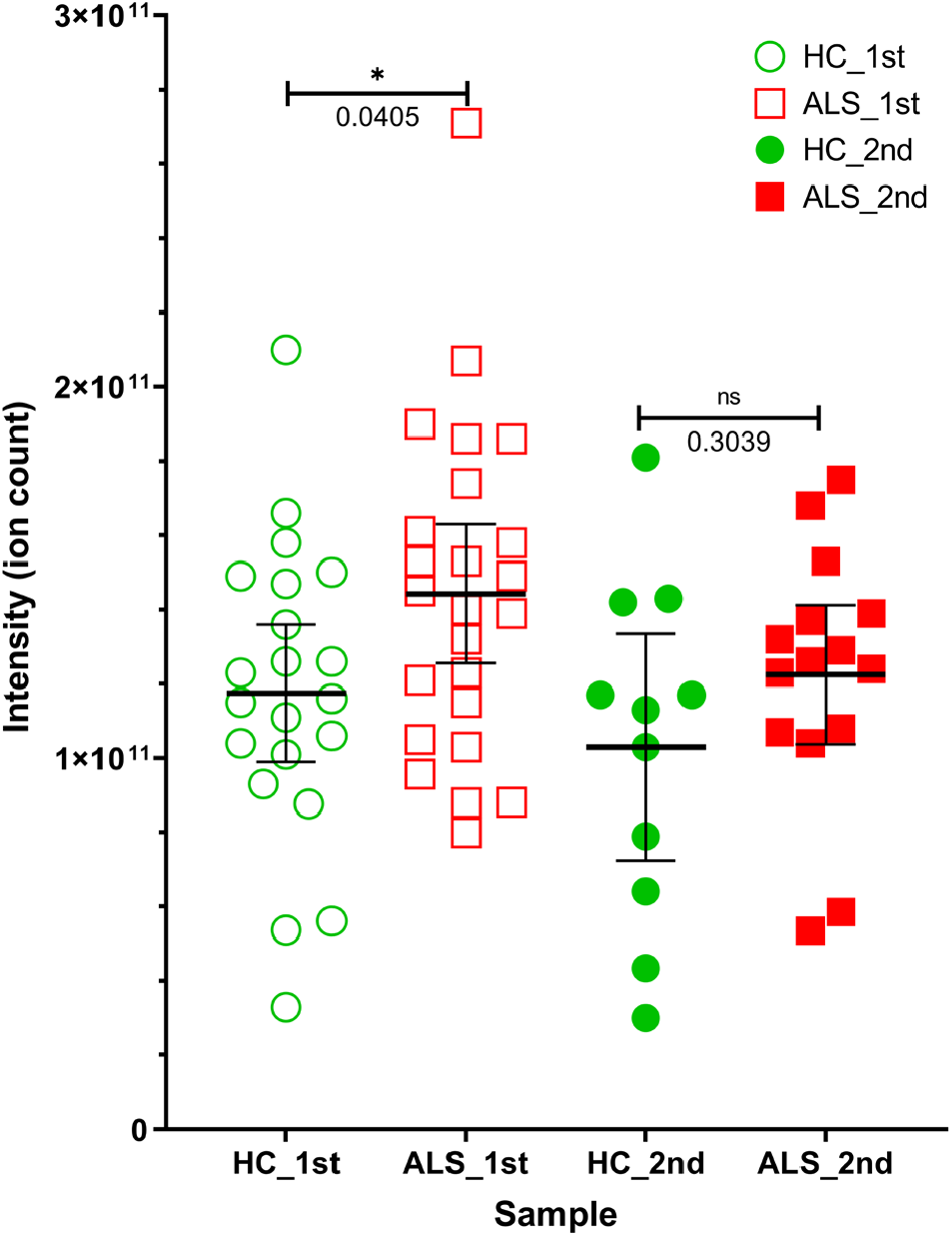
Total intensity of reversibly oxidised Cys (proteins) in HC and ALS groups at first and second visits. The total intensity is the sum of all peptide intensities bearing reversibly oxidised Cys per sample. ns: not significant, *: p-value < 0.05, Welch’s t-test.

### Development of targeted MS-based glutathione oxidation assay

We developed a highly sensitive targeted nanoflow LC-MS/MS-based method (pseudo-MRM) to accurately measure GSH and tGSH in human CSF with high sensitivity (Figure 3). The GSSG concentration is determined from the difference between tGSH and GSH as the tGSH consists of free reduced GSH and newly formed reduced GSH generated from GSSG through reduction with Tris(2-carboxyethyl)phosphine (TCEP). The method employs an alkylation step using NEM to derivatise GSH into GS-NEM, which is more stable for storage and enhances the sensitivity of detection of this derivatised compound by MS. A heavy stable isotope labelled GS*-NEM was used as an internal standard in the processed samples to achieve accurate quantitation and data normalisation. Before pseudo-MRM experiments were designed, full MS/MS scans of GS-NEM and GS*-NEM were acquired on an Orbitrap Elite system with a collision energy of 35 eV to obtain the MS/MS fragment profile, as shown in Figures S1 A and B. Of these fragment ions, only the highest abundance ions containing glycine-13C2, 15N were selected for the pseudo-MRM experiments. These signature ions were also annotated with chemical structures of GS-NEM and GS*-NEM, as shown in Figures S1 A and B. In LC-MS analysis, both GS-NEM and GS*-NEM co-eluted in a single peak (Figure S1). GS-NEM (433 m/z) and GS*-NEM (436 m/z) ions were fragmented by collision induced dissociation (CID), and selected fragment ions of 304 and 287 m/z were monitored in pairs with their corresponding parent ion 433 m/z for GS-NEM, and fragment ions of 307 and 290 m/z were monitored for internal standard ion 436 m/z (GS*-NEM) (Figure S2). Other pairs of ions were also monitored to ensure that alkylation and reduction reactions were completed. There were no other forms of GSSG and GSH detected, indicating that the alkylating and reducing steps were complete. A linear coefficient of 0.9969 was obtained for the GS-NEM standard curve, corresponding to GSH concentrations ranging from 0.13 to 26 μM (Figure 4A). A total of 140 CSF samples were analysed using this MS method, and internal standards were run after every 20 samples to evaluate the performance of the nano-HPLC MS/MS system. As a result, a coefficient variation (CV) of 5.0% was obtained for the internal standards alone (Figure 4B), suggesting a good performance of the instrument system during MS runs.

**Figure 3:**
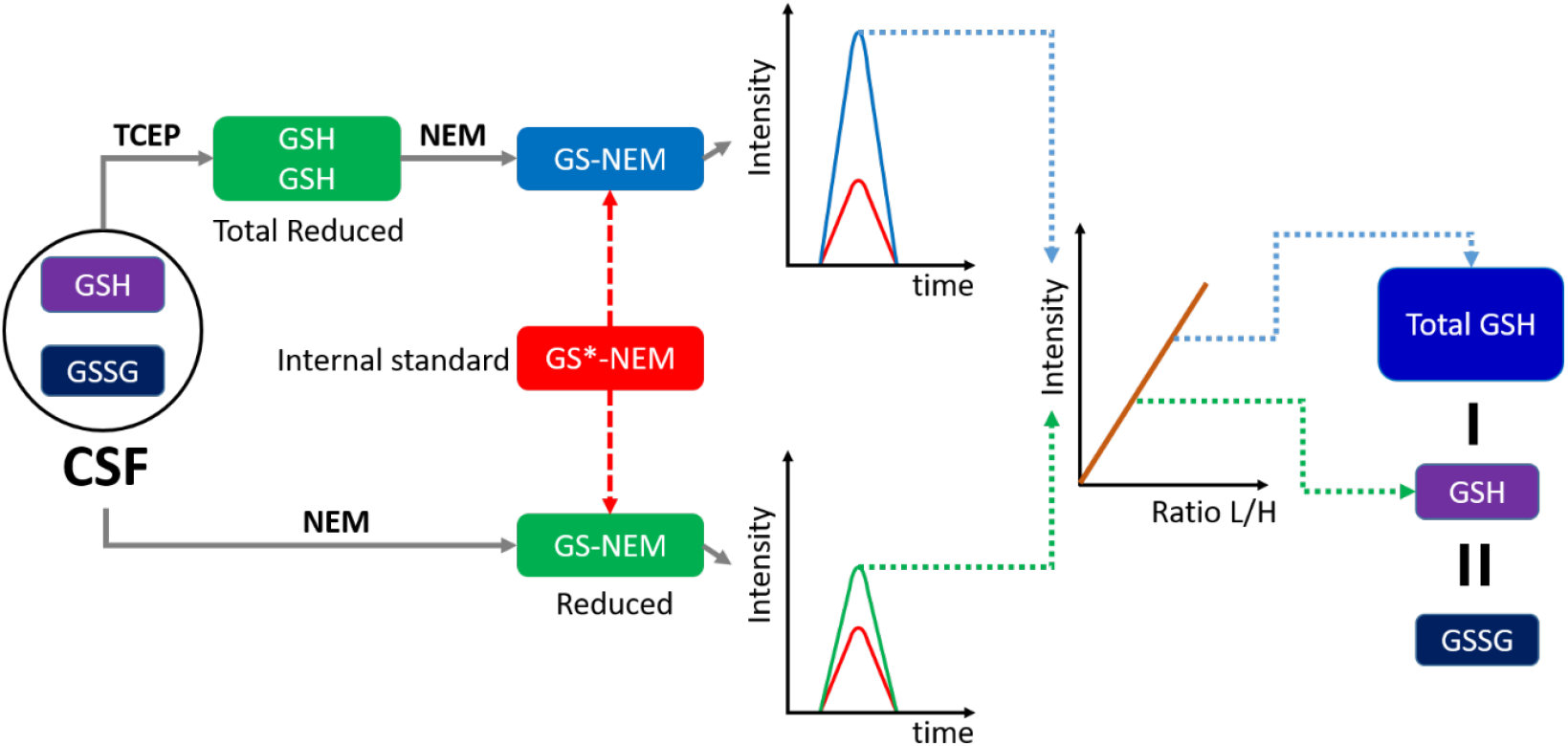
A workflow used to analyse glutathione oxidation in CSF samples. While GSH was directly alkylated with NEM, GSSG was first reduced using TCEP before being alkylated with NEM. A stable isotope-labelled internal standard GS*-NEM was added to each sample before MS analysis. GSH and Total GSH are measured in separate LC-MRM assays and levels of oxidised glutathione (GSSG) are inferred from the difference between GSH and Total GSH levels.

**Figure 4.**
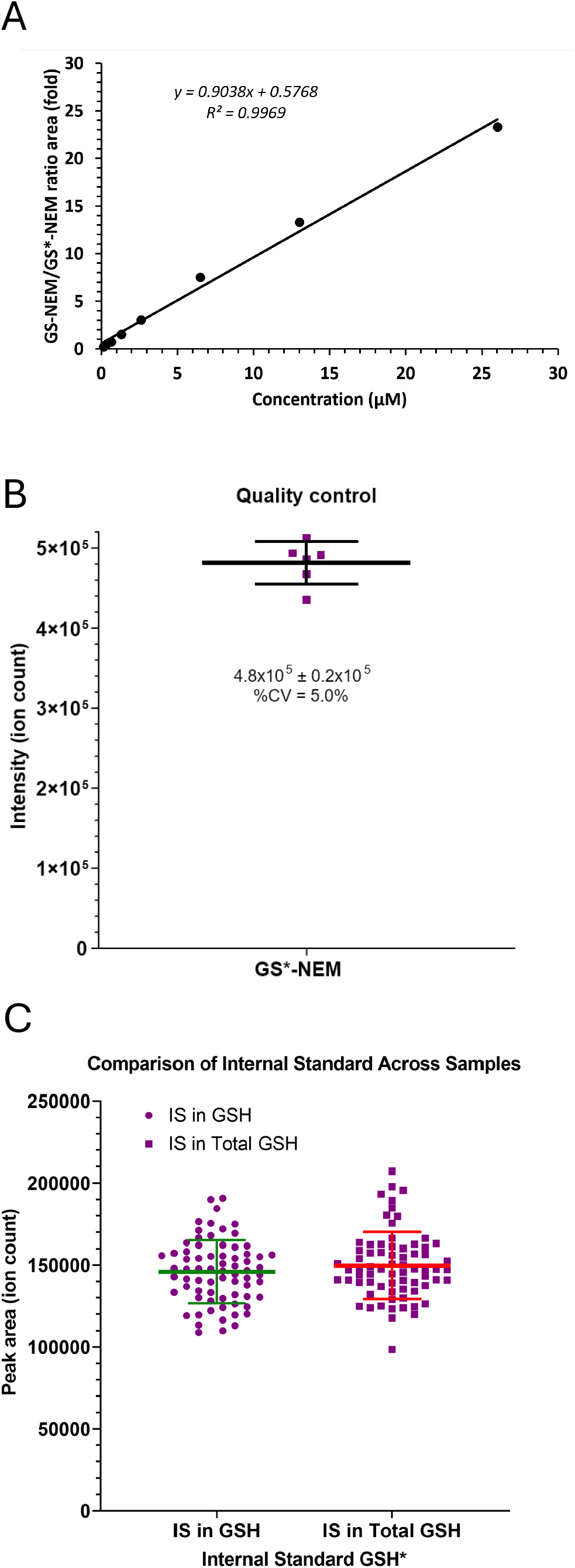
Performance of a targeted MS assay for glutathione oxidation. (A) The standard curve used to determine concentrations of GS-NEM and tGS-NEM in CSF. Intensity of heavy isotope labelled standard (IS) alone (B) and in CSF samples (C). IS was used for quality control of measurements and normalisation of data.

Signals of these internal standards were extracted and plotted in Figure 4C for two separate batches, GS-NEM and tGS-NEM measurements. CVs of internal standards in both batches were similar at 13.13 and 13.57% for reduced and oxidised measurements, respectively. The lower limit of detection (LLOD) and lower limit of quantitation (LLOQ) of GS-NEM (and also for tGS-NEM) were 0.04 and 0.13 μM, respectively, which were ten times lower than previously reported values [21] (0.4 μM and 1.5 μM for LLOD and LLOQ respectively). This suggests that our optimised MRM assay gives better detection sensitivity for the quantitation of glutathione in human CSF. Supplementary Table S4 shows the LLOD and LLOQ values in this study compared to other published values obtained by different approaches in various biological samples. Furthermore, CVs of 3.75 %, 4.91% and 5.1% were also determined for intraday, interday and accuracy for recovery tests, respectively.

### Oxidised glutathione is significantly elevated in the CSF of ALS patients

Concentrations of GSH, tGSH and GSSG in CSF were determined by GS-NEM and tGS-NEM measurements, and the detailed results are shown in Figure 4 (A and B for participants who attended the first and second visits, respectively, C for the ratio of GSSG/GSH). Data were obtained from 24 ALS patients (10 female, 14 male; aged 61 ± 10) and 20 healthy controls (13 female, 7 male; aged 59 ± 10). A summary of the mean group concentrations of GSH, tGSH and GSSG, and the GSSG/GSH ratio are also shown in Table 3. There were no significant differences in GSH concentrations between ALS and HC groups at the first or second visits or between visits (Figure 5 and Table 3). However, the concentration of tGSH in the ALS group was significantly higher than that of the HC group for the first (1.33-fold, p = 0.0215) and second (1.50-fold, p = 0.0143) visits. Similarly, the concentration of GSSG was also significantly increased in CSF of ALS compared to HC groups for both first (1.54-fold, p = 0.0041) and second (2.0-fold, p = 0.0018) visits. The GSSG/GSH ratio for the ALS groups was also significantly higher than those in the HC group at the second visit (2.84 vs 1.33, p = 0.0120). The mean concentrations of both tGSH and GSSG in CSF of ALS patients were almost unchanged between the first and second visits (0.44 mM ± 0.2 (1st) vs 0.43 mM ± 0.16 (2nd) or tGSH, and 0.18 mM ± 0.1 (1st) vs 0.18 mM ± 0.08 (2nd) for GSSG).

**Figure 5:**
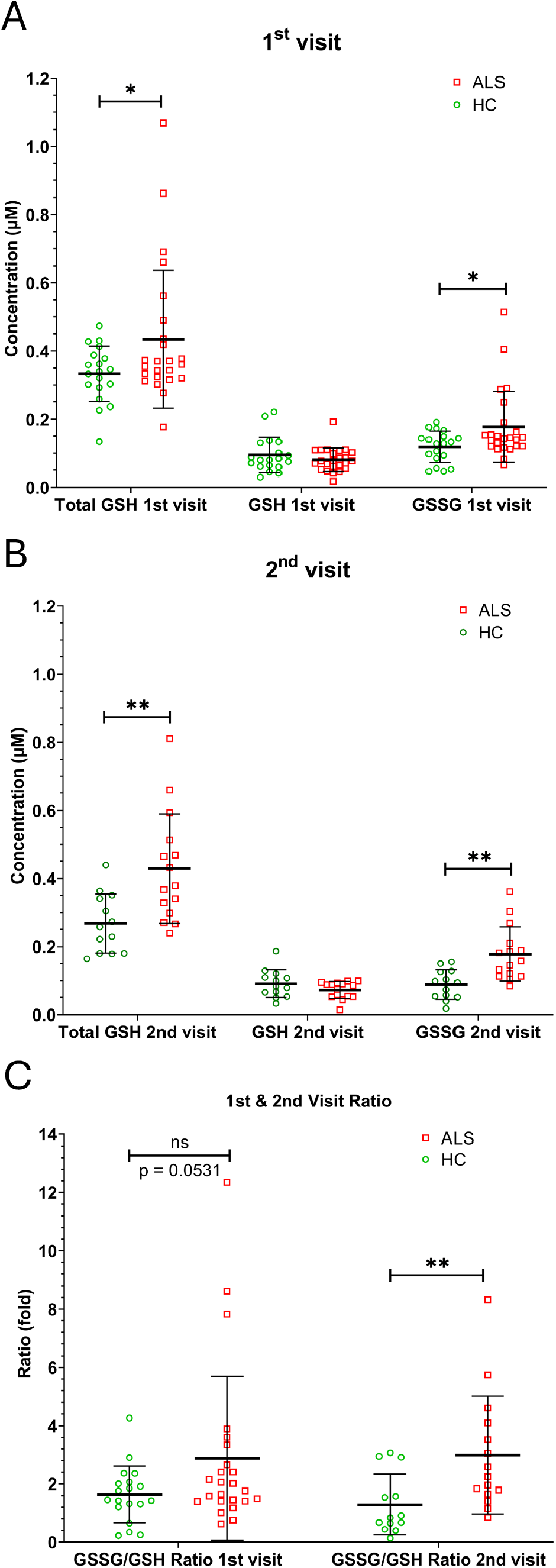
Concentrations of GSH, tGSH, GSSG and GSSG/GSH ratio (C) in ALS and HC CSF groups for the first (A) and second (B) visits. Green circles and red squares represent healthy controls and ALS cases, respectively; ns: not significant, *: p-value < 0.05, ** p-value < 0.01, Welch’s t-test.

### Biomarker correlations with clinical parameters

We used Pearson correlation and simple linear regression models to identify correlations between potential biomarkers and clinical parameters of ALS patients (Supplementary Table S4). Among proteins listed in Table 2, there were four proteins at the first visit and two proteins at the second visit of ALS patients that positively correlated with the disease progression rate. These included: (first visit) complement factor D (CFD) (p = 0.0436), complement factor H (CFH) (p = 0.0330), low-affinity immunoglobulin gamma Fc region receptor III-A (FCGR3A) (p = 0.0021), and alpha-1-antichymotrypsin (SERPINA3) (p = 0.0051); (second visit) pyruvate kinase (PKM) (p = 0.0344) and chitotriosidase-1 (CHIT1) (p = 0.0134). Among these proteins, the protein SERPINA3 also negatively correlated with the ALSFRS-R score (p = 0.0444) at the first visit of ALS patients. Finally, neurosecretory protein (VGF) showed a positive correlation with symptom onset (p = 0.0284). There was also a positive correlation between the increased progression rate in ALS patients (n = 7) between two visits (4 months apart) and the increased expression of Carbonyl reductase [NADPH] 1 protein (CA11) (p = 0.0069). In contrast, there was a negative correlation between the increased ALSFRS-R score and the changed expression of Phosphoglycerate kinase 1 (PGK1) (p = 0.0407).

Both tGSH and GSSG concentrations correlated positively with disease duration until the first visit, and the GSSG/GSH ratio also showed a positive correlation with disease duration until the first sample date. Linear regression models were determined for tGSH, GSSG and GSSG/GSH ratio, respectively, as follows: Y = 0.006783*X + 0.3122 (F = 9.581, p = 0.0055), Y = 0.003409*X + 0.1150 (F = 8.901, p = 0.0071) and Y = 0.07999*X + 1.409 (F = 6.042, p = 0.0227). Where X is disease duration until the first sample date (in months), and Y is tGSH (μM), GSSG (μM), or GSSG/GSH ratio (fold), respectively. Although not many reversibly oxidised protein Cys sites were significantly different in their abundance between ALS and HC groups, there were positive correlations between the total intensity of reversibly oxidised Cys and the ratio of GSSG/GSH in ALS patients for both visits, (p = 0.0193, p = 0.0367 for the first and second visits, respectively). This reflects the simultaneously increased level of reversible Cys oxidation of proteins and increased GSSG/GSH ratio in ALS patients. Furthermore, this correlation is significantly increased between two visits (4 months apart) since there was a positive correlation found between these time points (p = 0.0405) (Supplementary Table S4).

## Discussion

The goal of this study was to develop an approach to measure biomarkers of disease progression relevant to oxidative stress that could be used for target engagement studies in clinical trials. Therefore, we developed a workflow using MS-based approaches to measure protein abundance, cysteine oxidation, and the abundance and oxidation state of glutathione in CSF from a cohort of patients with ALS and a group of healthy controls at two time points. To reduce the very high dynamic range of protein abundance in CSF, we immunodepleted samples to remove the top 14 most abundant proteins. This enhanced the proteome coverage up to 3.5-fold compared to non-depleted CSF (data not shown). As a result, 1,561 proteins were detected in the entire data set, and 699 proteins were quantified in 70% of replicates (for each group, HC 1st, HC 2nd, ALS 1st, ALS 2nd).

We identified several proteins with significantly increased abundance in ALS compared to the HC group in both consecutive visits. These included chitinase-3-like protein 1 and 2 (CHI3L1 and CHI3L2), chitotriosidase-1 (CHIT1) and collagen alpha-1(XVIII) chain (COL18A1), and our data confirm these proteins as candidate biomarkers of ALS as previously reported [21] [46]. So far, CHI3L1 and CHIT1 in CSF have been identified as potential biomarkers in Alzheimer’s disease (AD), multiple sclerosis (MS) and other neurological diseases [22]. The levels of CHIT1, CHI3L1 and CHI3L2 in CSF of ALS patients were reported to be elevated compared to HC groups (using ELISA) [23, 24]. Furthermore, CHIT1 and CHI3L2 correlated with disease progression rate, and CHI3L1 correlated with the degree of cognitive dysfunction. CHIT1 levels were also associated with survival in multivariate models, and chitinase levels were longitudinally stable [23, 24]. However, in our data, the abundance of CHIT1, CHI3L1, CHI3L2, and collagen alpha-1(XVIII) chain did not change between the two visits in the ALS group, indicating that their expression level was stable as the disease progressed.

Protein alpha-1-antichymotrypsin (SERPINA3) was significantly elevated in ALS patients (compared to the HC group) at the first visit, suggesting it could play a potential role in ALS development since it positively and negatively correlated with disease progression rate and ALSFRS-R score, respectively. Moreover, the increased expression of alpha-1-antichymotrypsin together with CFH, C7, MASP1, SERPING1, FCGR3A, CFD, CHI3L1 and CHI3L2 might involve the immune system response and response to stress as an imbalance of the immune response could contribute to excessive inflammation occurring early during disease progression [25].

In terms of disease progression, there was a significant positive correlation between the increase in protein CA11 expression and the increase in disease progression rate over time (4 months apart). This means that the increase in progression rate from the first to the second visit is positively correlated with the increased abundance of protein C11 between these time points in ALS patients. The protein CA11 is a member of the carbonyl reductase family of enzymes, which play a role in the reduction of various carbonyl compounds, including reactive oxygen species (ROS) and oxidative stress. This protein has a potential role in neuroprotection as it is essential for neuronal cell survival and protection against oxidative stress [26]. Additionally, the carbonic anhydrase family has been shown to have a role in the regulation of the immune system, including the activation of microglia [27].

We next investigated if any oxidative biomarkers could be identified in the CSF of ALS patients and found that several proteins exhibited significantly altered cysteine oxidation in ALS versus the HC group (Supplementary Table 1). Brevican core protein (BCAN), exhibited increased (reversible) oxidation at Cys688 and Cys699 in ALS vs HC groups in both consecutive visits. This protein is a CNS-specific extracellular matrix proteoglycan and is degraded by extracellular metalloproteinases, proposing an unknown transport mechanism from the brain parenchyma into CSF [28]. Protein dipeptidyl peptise 2 DPP7 contained two redox cysteine sites (Cys332 and Cys338) more oxidised in ALS than HC for both visits. This protein is a member of the serine peptidase and plays a key role in the degradation of oligopeptides [29]. Proline-specific dipeptidyl peptidases (DPPs) have emerged as targets for drug development. Cys332 and Cys382 form a disulphide bond in the loop insertion, defining dipeptidyl aminopeptidase specificity and acting to stabilise the long loop [30]. Our data could indicate that more disulphide bonding occurs in ALS patients than in the HC group. Although 17 cysteines (from 14 different proteins) were irreversibly oxidised (O2 and O3), none of these were significantly altered in abundance in ALS compared to HC groups, but some proteins increased their irreversible oxidation in both ALS and HC between the first and second visits (Supplementary Table S2).

We next tested whether the level of glutathione or its oxidation state was altered in ALS versus healthy controls and between consecutive visits in the ALS cohort. The LC-MRM approach has been widely used in targeted metabolomics and proteomics analyses because of its high specificity, selectivity, accuracy, precision and robustness. Several MRM methods were successfully developed to measure glutathione levels in blood [31] and other biological samples using different MS-based approaches [32-34] (Supplementary Table S3). However, to our knowledge, there are no previous reports using this approach to measure glutathione in the CSF, which has much lower levels of glutathione compared to other biofluids (see Supplementary Table S3) and, therefore requires the higher sensitivity afforded by nano-flow chromatography.

In our MRM method development process, we generated a GSH standard curve for calculating GSH and tGSH concentrations in human CSF. Compared with the GSH standard curves reported [31] and [35], the slope of our standard curve was consistent with their standard curves, 0.9038 in our study compared to the slope of 0.9177 and 0.932 in these reports, respectively. A linear coefficient R2 = 0.9969 in our data was in line with R2 = 0.999 and R2 = 0.9986 found in [31] and [35], respectively. Furthermore, a linear dynamic range (0.13 – 26 μM) in our data is much lower than in these previous reports (25 – 500 μM and 8 – 256 μM as reported in [31] and [35], respectively). This allowed us to successfully measure the much lower glutathione levels in CSF compared to blood.

When measuring glutathione (GSH and GSSG) in clinical samples, it is important to inhibit the auto-oxidation of GSH (into the GSSG form), which could occur during sample collection and preparation. Therefore, we utilised NEM to block the sulfhydryl group on the cysteine residue of GSH via alkylation (to form derivatised GS-NEM) in order to prevent GSH auto-oxidation [36]. This also helps to prevent the enzymatic reduction of GSSG and inhibits glutathione reductase activity [36]. The concentrations of tGSH detected in the HC group (0.33 ± 0.08 and 0.28 ± 0.08 μM for the first and second visits, respectively) were similar to those previously reported in which tGSH was measured by a spectrophotometric approach [37]. It is noted that only healthy males participated (n = 26) in that study. The mean tGSH concentrations of 0.12 ± 0.02, 0.18 ± 0.01 and 0.14 ± 0.01 μM were reported for three different fractions 0–6, 7–12 and 13– 18 mL, respectively [37], while our data from a male healthy volunteer group for the first and second visits were 0.31 ± 0.10 (n = 7) and 0.29 ± 0.09 μM (n =5), respectively. The concentrations measured in our study were approximately 40% higher than the values previously reported [37]. This difference could result from the different methodologies used. However, the difference, in our opinion, was not significant, and we considered that our data did not conflict with the levels reported in the literature. Subsequently, this allowed us to examine if we could use levels of glutathione and/or its oxidation as potential biomarkers of oxidative stress in ALS.

Oxidative stress plays a key role in the pathophysiology of ALS [38, 39], but most antioxidants, including glutathione, have failed to slow ALS progression in clinical trials [40, 41]. Several potential biofluid-based biomarkers have been discovered for ALS [42-44], including neurofilament light chain (NfL), which is considered the best-performing candidate for therapeutic trials [45]. Unfortunately, neurofilament light chain was not detected in our data, potentially due to relatively low abundance in CSF or loss of the protein during the immunodepletion step. The failure to detect neurofilament light chain has been reported in other global proteomic studies of CSF from ALS patients [46] but has been successfully detected using immunoprecipitation– tandem mass spectrometry method in which peptides derived from the protein are specifically enriched from CSF after digestion, before MS analysis [47].

Our data show that three crucial parameters, including tGSH and GSSG concentrations in ALS, were significantly higher than those in the HC group at the first visit and tGSH and GSSG concentrations and the GSSG/GSH ratio in ALS at the second visit (Table 3 and Figures 4B and C). Compared to the healthy control group, the mean value of each parameter (tGSH, GSSG, and GSSG/GSH ratio) was almost unchanged in the ALS group for the second compared to the first visits (4 months apart), suggesting that the levels of these molecules remained high and unchanged once ALS is established. Our findings support the proposal that either tGSH or GSSG could be used as potential biomarkers in ALS.

There were strong positive correlations between disease duration until the first sample date and tGSH, GSSG and GSSG/GSH ratio parameters in the CSF of ALS patients for disease duration until the first sample date vs tGSH, GSSG and GSSG/GSH ratio, respectively. This suggests an accumulation of oxidative stress biomarkers over time, though these glutathione derivatives were relatively stable over the 4-month interval between sample collections in this study. There was no difference in the concentrations of GSH, tGSH and GSSG between slow-, intermediate-, and fast-progressing ALS cases measured by the rate of change of the ALS-FRS-R score per month.

The high concentrations of tGSH, GSSS, and GSSG/GSH ratio, as well as total reversibly oxidised Cys content, reflected the imbalance between the antioxidant defence system and ROS production, contributing to the development and progression of motor neuron injury in patients with ALS [48]. The measurement of tGSH, GSH and corresponding GSSG/GSH ratio could be used as biomarkers in ALS as the GSSG/GSH ratio showed a strongly positive correlation with the level of Cys oxidation in ALS patients, and the longer the disease duration, the stronger was the correlation observed. A higher GSSG/GSH ratio was also reported in a SOD1-ALS cell model [48] and a SOD1-ALS mouse model [49].. Our data, especially the positive correlation between the protein Cys oxidation level and GSSG/GSH ratio, could offer potential biomarkers for the evaluation of oxidative stress mechanisms contributing to neurodegeneration in ALS and the effects of therapeutic interventions targeting oxidative stress.

## Conclusion

We have employed various MS-based approaches to simultaneously measure protein abundance and protein oxidation in the CSF of ALS patients compared to healthy controls. We identified several proteins with significantly altered abundance in ALS and cysteine-based oxidative proteomics revealed a global increase in protein oxidation in ALS patients. In our workflow for human CSF sample preparation, we utilised a low molecular weight fraction, a by-product of immunodepletion, for the targeted measurement of glutathione oxidation using a sensitive nano-flow pseudo-MRM approach. Using this method, we measured glutathione levels in the CSF of both ALS patients and HC volunteers in longitudinal samples over a four-month period. We found the concentrations of tGSH and GSSG and the corresponding GSSG/GSH ratio were significantly elevated in the ALS compared to the HC group. Furthermore, there was a positive correlation between disease duration at the time of the first CSF sample with tGSH, GSSG and the GSSG/GSH ratio, reflecting that the longer the ALS disease duration, the higher the levels of tGSH or GSSG. These results support further evaluation of tGSH and GSSG in the setting of therapeutic intervention in clinical trials of therapies targeting oxidative stress.

## Acknowledgements

This work was supported by a Medical Research Council award (COEN-Centres of Excellence for Neurodegeneration MRC/S004920/1) to PJS, RJM and MC and by the ALS Association AMBRoSIA (A multi-centre biomarker resource resources strategy for ALS) programme (Oct15/972-797) to PJS, MRT and AM. The study was also supported by the NIHR Sheffield Biomedical Research Centre (NIHR203321) and the NIHR Sheffield Clinical Research Facility. Grateful thanks are extended to those ALS patients and healthy volunteers who generously donated biosamples for research.

## Materials and Methods

### Chemicals

Iodoacetamide (IAM), Dithiothreitol (DTT) and N-Ethylmaleimide (NEM) (Merck, UK) were dissolved in water to make a stock of 0.25 M, while TCEP 0.5M (Merck, UK) was used as a stock. Both GSH and GSSG were purchased from Merck, while heavy isotope labelled GSH (glycine-13C2, 15N-Glutathione) (GSH*) was purchased from Cambridge Isotope (UK). All of these reagents were resuspended in water to make a stock of 1 mg/mL for each and used as standards and internal standards (GSH*). HPLC-MS grade water, trifluoroacetic acid (TFA), formic acid (FA), acetonitrile (ACN) and 3kDa Amicon ultra-0.5 centrifugal filter unit were also purchased from Merck (UK), while spinning C18 cartridges were purchased from Thermo (UK).

### CSF samples

Samples were collected from participants enrolled in the observational ‘NRF2 Biomarkers’ study. The study was approved by an NHS Research Ethics committee (reference: 18/Y.H./0253). Patients were recruited at the Royal Hallamshire Hospital, Sheffield, UK, and provided written informed consent. A lumbar puncture was performed, together with clinical assessments and characterisation of the disease state and rate of disease progression. CSF samples were immediately placed on ice and centrifuged at 3500 rpm for 10 min at 4°C within 1 h of sampling and then transferred to a –80°c freezer. They were subsequently stored in liquid nitrogen. Demographic and clinical details of the subjects are shown in Table 1.

#### Protein sample preparation and immunodepletion

A protease inhibitor cocktail (leupeptin 1 μM, bestatin 130 μM and aprotinin 0.15 μM, all in working concentration) was added to 800 μL of CSF. Samples were concentrated using amicon ultra-15 centrifugal filter units (Millipore) with a 3 kDa molecular weight cut off (MWCO) at 4oC for 90 min at 4,000 x g. After centrifugation, the concentrated protein (∼ 200 μL) was collected and transferred to a new 1.5 mL Eppendorf tube. The flow-through fraction (< 3 kDa) was collected and used for glutathione measurement described below. Concentrated protein samples were then immunodepleted using a multi-affinity removal spin cartridge human-14 (0.45 mL, Agilent Technologies, UK) to remove high abundance proteins according to the manufacturer’s protocol with some modifications. Briefly, 200 μL of buffer A was added to the concentrated CSF sample and incubated in the cartridge at room temperature for 5 minutes before centrifuging at 100 x g for 1.5 minutes. The supernatant containing low abundance proteins was collected into a new clean 2 mL Eppendorf tube. Another 400 μL of buffer A was added into the cartridge to elute unbound (low abundance) proteins. The supernatant was then collected by centrifugation at room temperature for 3 min and then combined with the previous supernatant. The combined low abundance protein fraction was transferred to another centrifugal filter unit (3kDa MWCO) and centrifuged at 4 oC for 90 min at 4,000 x g to concentrate proteins. Subsequently, the concentrated proteins (∼200 μL) inside the filter unit were transferred to a new 2 mL Eppendorf tube. Then 1mL of (−20oC) ice-cold acetone was added into the tube and left at –20oC overnight to precipitate protein. The pellet of precipitated protein was collected by centrifugation at 21,000 x g for 15 min at 4oC. Acetone was then discarded, and the pellet was left to dry before 25 μL of protein extraction buffer (5% sodium dodecyl sulfate (SDS) in 50 mM triethylammonium bicarbonate TEAB pH 7.1 was added to dissolve the protein pellet. A volume of 4 μL was used for total protein concentration determination using a microBCA assay (Thermo, UK). 25 μg of protein from each sample was then digested in a suspension trapping device (S-trap) (Protifi, USA). Proteins were firstly alkylated using 15 mM iodoacetamide (IAM), shaking at 850 rpm in a thermomixer (Eppendorf, UK) in the dark for 45 min at room temperature, and then reduced using 35 mM DTT with the same previous condition before subjecting to a second alkylation by 60 mM NEM using the same conditions. Subsequently, samples were centrifuged at 21,000 x g for 15 min before transferring to an S-Trap microcolumn (≤ 100 μg) (Protifi, USA) for protein digestion according to the manufacturer’s protocol. Briefly, proteins were trapped twice and washed 3 times with 100 mM TEAB pH 7.5 in 90% methanol before digestion in the S-trap using trypsin (Pierce, UK) (1:10 ratio of trypsin: protein) at 37oC overnight. Peptides were subsequently eluted with (60 μL each) 50 mM TEAB pH 8.5, and 0.2% formic acid (FA) then twice with 0.5% FA in 50% acetonitrile. The eluted peptides were dried in a vacuum concentrator (Eppendorf, UK) and resuspended in 50 μL of 0.5% FA for mass spectrometry analysis (MS).

### GSH preparation, reduction and alkylation

A workflow for processing and measuring glutathione in CSF is shown in Figure 2. For measurement of GSH in CSF, 300 μL of flow-through from the initial amicon concentration step above (< 3 kDa MWCO) was derivatised (alkylated) with NEM 20 mM and incubated in a thermomixer (Eppendorf, UK) at 25oC for 45 min at 850 rpm in the dark. For tGSH measurement, another 300 μL of flow-through CSF was first reduced using 10 mM TCEP at 25oC for 45 min at 850 rpm and then derivatised with 20 mM NEM. To clean up derivatised samples (GS-NEM), all samples were subjected to solid-phase extraction using C18 cartridges. TFA was added to the derivatised samples to reach a final concentration of 0.1%. The samples were then centrifuged at 21,000 x g for 15 min; supernatants were collected for desalting, which was performed according to the manufacturer’s protocol with some modifications. Briefly, the binding step was performed twice to maximise the binding of GS-NEM to C18 material; the C18 columns were washed three times with 300 μL of 0.1% TFA and then eluted with 25% ACN in 0.1% TFA and collected into a 2 mL Eppendorf tube. All samples were then dried in a vacuum concentrator (Eppendorf, UK) and stored at –80oC for MS-based analysis. Both light (GSH) and heavy stable isotopes (GSH*) of glutathione were also derivatised with NEM in the same way described for the CSF samples. Furthermore, to evaluate both the efficiency of the reduction step and the loss of GS-NEM during the C18 cleaning step, a known amount of SGGG (1.3 μM) was used for reducing, alkylating and subjecting to the desalting step and treated the same way compared to CSF samples.

#### GSH standard curve and MS performance evaluation

A linear standard curve of GS-NEM, in which 4 μL of 1.3 μM GS*-NEM was added into 38 μL of 0.5% FA for each GS-NEM concentration to serve as an internal standard. The standard curve was performed and established with concentrations of GS-NEM consisting of 0.13, 0.33, 0.65, 1.30, 2.60, 6.51, 13.02 and 26.03 μM. Triplicate measurements were madce for each GS-NEM concentration. Quality control during MS analysis was also performed using the internal GS*-NEM standard to check the accuracy of glutathione measurements and the MS performance. A volume of 18 μL, in which every 4 μL of GS*-NEM (1.3 μM) was added to 38 μL of 0.5% FA, was run on an Orbitrap Elite (Thermo, UK) every 24h. The intensities of these quality control samples were then analysed to evaluate the performance of the MS instrument. Furthermore, a blank was also run after every 6 CSF samples to monitor for potential chromatography contamination. The recovery of GSH during sample preparation was assessed by spiking 50 fmol of GSH into CSF samples (n = 3) before sample preparation, and CSF control samples (without the addition of GSH) (n = 3) were also performed. The recovery of GSH was calculated as follows: Recovery (%)= (Amount of measured GSH amount in spiked sample-amount of measured GSH in control sample)/(Theoretical amount of spiked GSH) X 100

### Nano-HPLC and MS analysis

#### Oxi-proteomics analysis

18 μL of resuspended peptides was injected and analysed by nanoflow LC-MS/MS using an Orbitrap Elite hybrid mass spectrometer (Thermo, UK) equipped with a nanospray source, coupled to an Ultimate RSLCnano LC System (Dionex, UK) and Tune Plus for Orbitrap Elite (Thermo). Peptides were desalted on-line using a nano C18 trap column, 75 μm ID x 20 mm (Thermo) and separated using a 130-min gradient starting from 3 to 40% buffer B consisting of 0.5% FA in 80% ACN on an EASY-Spray column, 50 cm × 50 μm ID, PepMap C18, 2 μm particles, 100 Å pore size (Thermo). The Orbitrap Elite was operated with a cycle of one MS (in the Orbitrap) acquired at a resolution of 120,000 at m/z 400, with the top 20 most abundant multiply charged (2+ and higher) ions in a given chromatographic window subjected to MS/MS fragmentation in the linear ion trap. An FTMS target value of 1e6 and an ion trap MSn target value of 1e4 were used with the lock mass (445.120025) enabled. Maximum FTMS scan accumulation time of 200 ms and maximum ion trap MSn scan accumulation time of 50 ms were used. Dynamic exclusion was enabled with a repeat duration of 45 s with an exclusion list of 500 and an exclusion duration of 30.

##### GSH MS analysis

The dried derivatised CSF samples were resuspended in 38 μL of 0.5% FA, and 4 μL of GS*-NEM (1.3 μM) was then added to serve as an internal standard. 18 μL of each sample was injected and analysed by a nano-HPLC (Dionex, UK) coupled to an Orbitrap Elite (Thermo, UK). Derivatised compounds were separated using a PepMap RSLC C18 column (2 μm, 100 Å, 50 μm x 15 cm) (Thermo, UK) operated at 40oC at a flow of 0.25 μL /min during a 35 min gradient generated by a solvent A consisting of 0.1% FA in water and solvent B containing 0.1% FA in 80% ACN. The gradient was run at 3% of solvent B for 10 min, then ramped up to 45% solvent B over 15 min and a second ramp to 90% solvent B over 2 min, isocratic run at 90% solvent B for 5 min, then decreased to 3% B for 5 min. The MS was activated 10 minutes after starting the HPLC and acquired data over 25 minutes. Both GS-NEM and GS*-NEM co-eluted at retention time (Rt) 19 min (on HPLC chromatography) or appeared at Rt = 9 min (on MS chromatography). GS-NEM, GS*-NEM and other analytes in each CSF sample were ionised using electrospray ionisation (ESI) operated at 50 oC, sheath gas 6, and spray voltage 4000 V, with a capillary temperature of 300 oC. The Orbitrap Elite MS was operated in positive ion trap mode with a collision energy of 35. For quantitative analysis of both GS-NEM and GS*-NEM, the following transitions were monitored using pseudo-MRM (multiple reaction monitoring) as follows: for GS-NEM (m/z): 433 ⟶ 304 and 433 ⟶ 287, for GS*-NEM (glycine-13C2, 15N-GS-NEM) (m/z): 436 ⟶ 307 and 436 ⟶ 290. Furthermore, un-derivatised GSH and intact GSSG forms were also monitored to ensure that both reduction and alkylation processes were complete. Therefore, transitions (m/z) of 308 ⟶ 179 and 308 ⟶ 162 (for monitoring GSH), and 307 ⟶ 299, 307 ⟶ 276, 613 ⟶ 484 and 613 ⟶ 355 (for monitoring GSSG in [M+2H]2+ and [M+H]1+ forms respectively). However, neither underivatised GSH or GSSG were detected in CSF samples. MS data were acquired with Xcalibur V 3.0.63 software (Thermo, UK).

### Data analysis

#### Oxiproteomics analysis

Raw MS data were processed with MaxQuant V.1.6.10.43, searching against a human UniProt sequence database downloaded from https://www.uniprot.org/ (Mar 2021) using the following search parameters: Trypsin/P for digestion with 2 missed cleavages, methionine oxidation (M), N-terminal protein acetylation, and Carboxyamidomethylation (CAM, 57 Da) and N-ethylmaleimide (NEM, 125 Da) on cysteine were set as variable modifications. The first and main searches were carried out with MS tolerance of 10 and 5 ppm, respectively. Label-free Quantification (LFQ) was enabled with a minimum ratio count of 2, a minimum number of neighbours of 3 and an average number of neighbours of 6. PSM and protein match thresholds were set at 0.1 ppm. A protein false discovery rate (FDR) of 0.01 and a peptide FDR of 0.01 were used for identification level cut-offs. LFQ was performed using only peptides containing no PTM (except for acetyl protein N-term). Peptides containing cysteine modified by CAM was considered as initially reduced Cys-containing peptides while peptides containing NEM modification on cysteine were oxidised Cys-containing peptides. The levels of both reduced and oxidised Cys-containing peptides were used to determine the levels of cysteine oxidation in ALS and HC groups. Changes in cysteine oxidation levels were normalised to protein expression determined by LFQ.

The peptide and protein data from Maxquant outputs were further analysed using Perseus software V.1.6.1.50. The ProteinGroup.txt file was used for LFQ, while IAMPeptide.txt and NEMPeptide.txt files were used for cysteine oxidation calculations. The reversed and potential contaminant proteins/peptides were removed before analysis. For Cys-containing peptide analysis, an expand site table step was performed before any analysis was carried out. Intensity of proteins/peptides was transformed to log2 before further analysis. At least 70% valid value for one of the group was used for data filtering, proteins in each sample were normalised to median and missing values were imputed using normal distribution with width of 0.3 and down shift of 1.8. Finally, multiple ANOVA tests were performed to investigate differently abundant proteins in ALS vs HC comparisons as well as between two visits to examine their association with disease progression.

The lists of differently significant expressed (oxi)proteins (Tables 2 and S2) were used for determination of correlations with clinical parameters using a simple regression model performed by Grapad.

### GSH analysis

MS data were analysed using Skyline version 21.1.0.218 using the small molecule analysis approach to obtain values of peak areas, and ratios of light (GS-NEM) and heavy (GS*-NEM) derivatised compounds. Ratios of light/heavy GS-NEM were referred to the standard curve to obtain concentrations of reduced and total reduced GSH. The obtained concentrations were then multiplied by a dilution factor of 7.14 (∼ 40 μL /300 μL) to calculate concentrations of GSH and tGSH in CSF. The concentration of GSSG was calculated based on a subtraction of tGSH and GSH concentrations. Extracted data were exported into data spreadsheets, and further statistical analyses were carried out using Excel 2016 (Microsoft, USA) and GraphPad Prism V 9.2.0 (GraphPad Software, USA).

Welch’s t-test was performed for each comparison to determine if any change of glutathione in CSF occurred between ALS and HC groups for 1st and 2nd visits. Using GraphPad, two-tailed Pearson correlation and a simple linear regression model were utilised to identify correlations between clinical parameters and glutathione in the CSF from ALS patients.

