## Supplemental Information for "Glutathione Oxidation in Cerebrospinal Fluid as a Biomarker of Oxidative Stress in Amyotrophic Lateral Sclerosis"

**Supplementary Information**

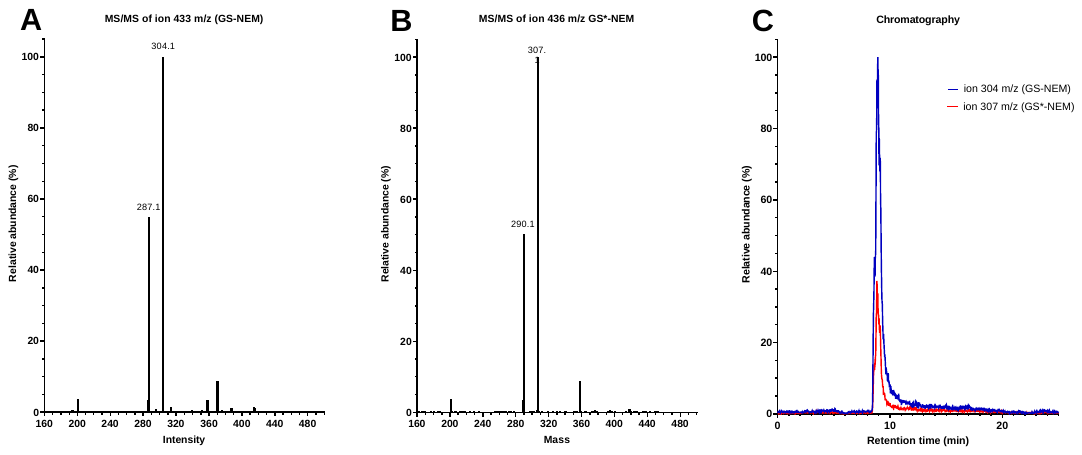

Figure S1. MS/MS fragments of GS-NEM (A) and GS*-NEM (B), and p-MRM chromatography of GS-NEM and GS*-NEM co-eluted from HPLC (C). Structures of selected ions used for p-MRM experiments are shown in Figs S2. A and B.

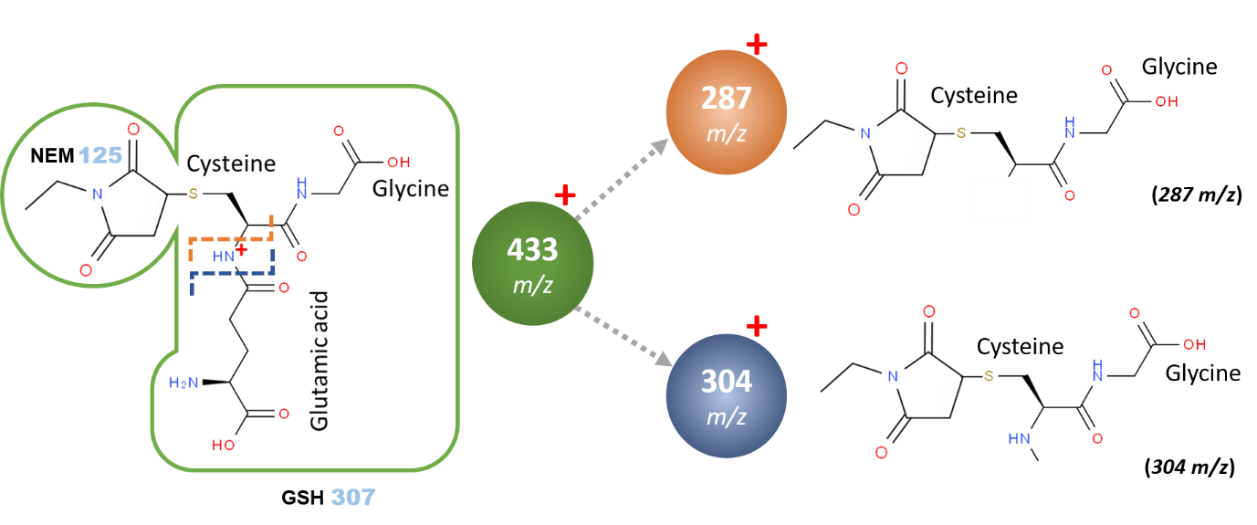

**A**

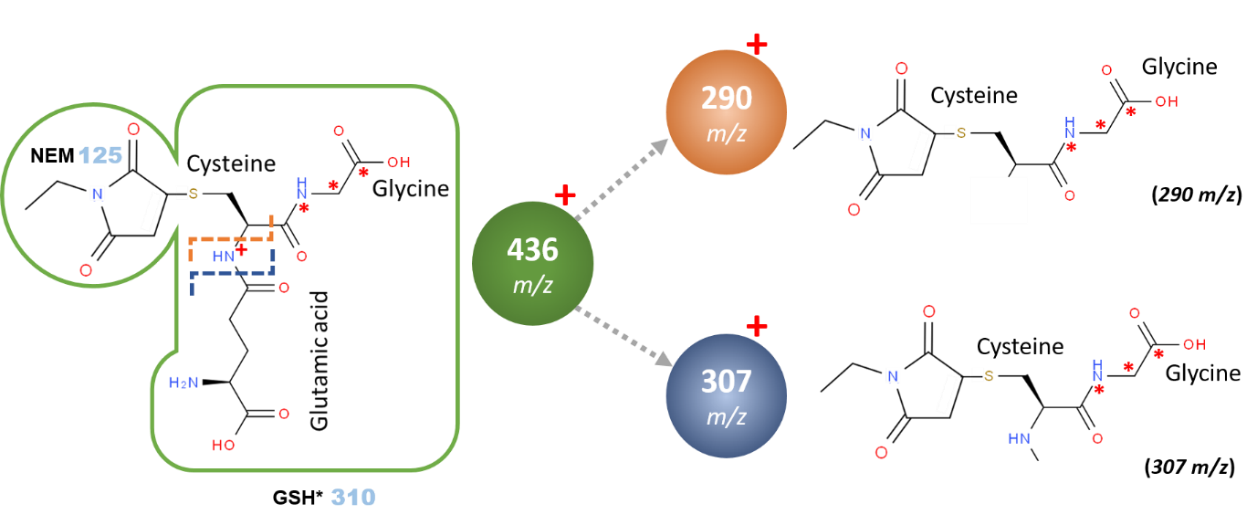

**B**

**Figure S2. Chemical structures of selected ions derived from the fragmentation of GS-NEM and GS*-NEM.**

**Supplementary Table S2.** List of oxi-proteins changed in their oxidative state on Cys residues.

*Note: Bold value indicates the fold change of a significantly different Cys oxidation site.*

| **AC** | **Name** | **Gene_Cys site** | **State** | **Fold ALS vs HC 1st** | **Fold ALS vs HC 2nd** |
| --- | --- | --- | --- | --- | --- |
| Q96GW7 | Brevican core protein | BCAN_C688 | Reversible | **1.96** | **2.93** |
| Q96GW7 | Brevican core protein | BCAN_C699 | Reversible | **1.21** | **2.39** |
| Q9UHC6 | Contactin-associated protein-like 2 | CNTNAP2_C558 | Reversible | 1.21 | **3.27** |
| G5E9G7 | Neurexin 2, isoform CRA_a | NRXN2_C299 | Reduced | 1.00 | **3.43** |
| P02748 | Complement component C9 | C9_C255 | Redox | 1.39 | **1.45** |
| Q9UHL4 | Dipeptidyl peptidase 2 | DPP7_C332 | Redox | **1.09** | **1.54** |
| Q9UHL4 | Dipeptidyl peptidase 2 | DPP7_C338 | Redox | **1.47** | **1.69** |
| B4E1Z4 | Complement Factor B | B4E1Z4_C564 | Redox | **2.26** | **3.56** |
| P01011 | Alpha-1-antichymotrypsin | SERPINA3_C261 | O3 | 2.84 | **6.92** |
| B4E1Z4 | Complement Fator B | B4E1Z4_C794 | O3 | 1.59 | ***-3.12*** |
| P41222 | Prostaglandin-H2 D-isomerase | PTGDS_C186 | O2 | **3.32** | -1.06 |

**Supplementary Table S3.** LLOD and LLOQ of GSH determined by various approaches and chemicals used for derivatisaion

| **Approach** | **Detection** | **Sample Type** | **Chemical** | **LLOD (µM)** | **LLOQ (µM)** | **Linear range/ regression model** | **Year** | **Reference** |
| --- | --- | --- | --- | --- | --- | --- | --- | --- |
| Enzymatic recycling | Absorbance at 405 nm | Rat liver/bile | M4VP |  | 0.375 | 0.38 ÷ 6.00 µM  Y = 0.000546*X  (r^2^ = 0.999) |  | [51] |
| HPLC-UV | UV detection at 330 nm | Blood (Erythrocytes) | DTNB | 0.041 | 0.135 | 0.5 ÷ 3.0 µM  (r^2^ > 0.99) |  | [52] |
| HPLC-Fluorescence | Fluorescence detection at 470/530 nm | Plasma | NBD-F | 0.03 | 0.10 | 0.1 ÷ 10 µM  Y = 37641*X + 5152  r^2^ = 0.9988 | 2012 | [53] |
| HPLC-MS/MS  (MRM) | MS | Whole blood | NEM | 0.4 | 1.5 | Y = 0.9177*X + 5.0443  r^2^ = 0.999 | 2013 | [21] |
| HPLC-UV |  | Cultured cells | NEM, DTT, mBrB |  |  |  | 2015 | [37] |
| HPLC-UV  HPLC-QTOF-MS | UV detection at 210 nm  MS | Cultured cells | NEM | 7.81 | 15.63 | 15.63 ÷ 1000  Y = 8.1448*X + 25.5998  r^2^ = 0.9997 | 2020 | [54] |
| HPLC-MS/MS  (MRM) | MS | Pig liver | NEM |  | 0.1 | 8 ÷ 256  Y = 0.0532*X + 0.0449  r^2^ = 0.9986 | 2014 | [55] |
| HPLC-MS/MS  MRM | MS | Blood | NEM and d_5_-NEM | 2 nM | 5 nM |  | 2018 | [56] |
| HPLC-MS/MS  MRM | MS | Whole blood, plasma and serum  Tissue  Cultured cells | NEM and d_5_-NEM |  |  | Y = 0.932*X + 2.49  r^2^ = 0.994 | 2020 | [35] |
| HPLC-MS/MS  Pseudo MRM | MS | CSF | NEM | 0.04 | 0.13 | 0.13 ÷ 26  Y = 0.9038*X + 0.5768  r^2^ = 0.9979 | 2023 | This study |

M4VP: 1-methyl-4-vinyl-pyridinium; DTNB: 5,50 -dithio-bis-(2-nitrobenzoic acid); NBD-F: 7-flouro-4-nitrobenzo-2-oxa-1,3-diazole; NEM: N-ethylmaleimide; mBrB: monobromobimane; DTT: dithiothreitol. LLOD and LLOQ reported in each study were converted to the same unit. Y: Detection signal, X: Concentration of GSH or (GS-NEM), µM.

**Supplementary Table S4**: Regression models displaying correlations between potential biomarkers and clinical parameters

| **AC** | **Protein name/Other** | **Gene/Other** | **Regression model** | **Where** | **Correlation with clinical parameter** | **F value** | **p value** |
| --- | --- | --- | --- | --- | --- | --- | --- |
| P00746 | Complement factor | CFD | Y = 0.1574*X + 3.078 | X: normalised intensity; Y: progression rate | Progression rate | 4.609 | 0.0436 |
| P08603 | Complement factor H | CFH | Y = 0.1345*X + 5.138 | X: normalised intensity; Y: progression rate | Progression rate | 5.209 | 0.033 |
| P08637 | Low affinity immunoglobulin gamma Fc region receptor III-A | FCGR3A | Y = 0.1987*X - 0.01980 | X: normalised intensity; Y: progression rate | Progression rate | 12.26 | 0.0021 |
| P01011 | Alpha-1-antichymotrypsin | SERPINA3 | Y = 0.2183*X + 7.728 | X: normalised intensity; Y: progression rate | Progression rate | 9.762 | 0.0051 |
| P14618 | Pyruvate kinase | PKM | Y = 0.5359*X + 3.566 | X: normalised intensity; Y: progression rate | Progression rate | 5.585 | 0.0344 |
| Q13231 | Chitotriosidase-1 | CHIT1 | Y = 6.207*X - 2.600 | X: normalised intensity; Y: progression rate | Progression rate | 8.188 | 0.0134 |
| P01011 | Alpha-1-antichymotrypsin | SERPINA3 | Y = -0.03806*X + 9.411 | X: normalised intensity; Y: ALSFRS-R score | ALSFRS-R score | 4.572 | 0.0444 |
| O15240 | Neurosecretory protein | VGF | Y = 0.09277*X - 4.866 | X: normalised intensity; Y: Symptom onset | Symptom onset | 8.243 | 0.0284 |
| P16152 | Carbonyl reductase [NADPH] 1 protein | CBR1 | Y = 6.948*X - 1.042 | X: Increased progression rate; Y: Protein abundance change | Increased progression rate | 19.54 | 0.0069 |
| P00558 | Phosphoglycerate kinase 1 | PGK1 | Y = -1.158*X - 0.3455 | X: Increased ALSFRS-R score; Y: Protein abundance change | Increased ALSFRS-R score | 7.516 | 0.0407 |
|  | Total reduced glutathione | tGSH | Y = 0.006783*X + 0.3122 | X: disease duration until first sample date; Y: tGSH (µM) | Disease duration until first sample date | 9.581 | 0.0055 |
|  | Oxidised glutathione | GSSG | Y = 0.003409*X + 0.1150 | X: is disease duration until first sample date; Y: GSSG (µM) | Disease duration until first sample date | 8.901 | 0.0071 |
|  | Ratio between oxidised and reduced glutathione | GSSG/GSH ratio | Y = 0.07999*X + 1.409 | X: disease duration until first sample date; Y: GSSG/GSH (fold) | Disease duration until first sample date | 6.042 | 0.0227 |
|  | Ratio between oxidised and reduced glutathione in the 1st visit | GSSG/GSH ratio | Y = 3.008e-11*X - 1.460 | X: the total intensity of reversibly oxidised Cys; Y: GSSG/GSH ratio |  | 6.376 | 0.0193 |
|  | Ratio between oxidised and reduced glutathione in the 2nd visit | GSSG/GSH ratio | Y = 3.577e-11*X - 1.272 | X: the total intensity of reversibly oxidised Cys; Y: GSSG/GSH ratio |  | 5.42 | 0.0367 |
